# Retinal ganglion cell vulnerability to pathogenic tau in Alzheimer’s disease

**DOI:** 10.1101/2024.09.17.613293

**Authors:** Miyah R. Davis, Edward Robinson, Yosef Koronyo, Elena Salobrar-Garcia, Altan Rentsendorj, Bhakta P. Gaire, Nazanin Mirzaei, Rakez Kayed, Alfredo A. Sadun, Alexander V. Ljubimov, Lon S. Schneider, Debra Hawes, Keith L. Black, Dieu-Trang Fuchs, Maya Koronyo-Hamaoui

**Author notes:** Corresponding author: Maya Koronyo-Hamaoui, PhD, Cedars-Sinai Medical Center, 127 S. San Vicente Blvd., A6212, Los Angeles, CA, USA 90048.

## Abstract

Accumulation of pathological tau isoforms, especially hyperphosphorylated tau at serine 396 (pS396-tau) and tau oligomers, has been demonstrated in the retinas of patients with mild cognitive impairment (MCI) and Alzheimer’s disease (AD). Previous studies have noted a decrease in retinal ganglion cells (RGCs) in AD patients, but the presence and impact of pathological tau isoforms in RGCs and RGC integrity, particularly in early AD stages, have not been explored. To investigate this, we examined retinal superior temporal cross-sections from 25 patients with MCI (due to AD) or AD dementia and 16 cognitively normal (CN) controls, matched for age and gender. We utilized the RGC marker ribonucleic acid binding protein with multiple splicing (RBPMS) and Nissl staining to assess neuronal density in the ganglion cell layer (GCL). Our study found that hypertrophic RGCs containing pS396-tau and T22-positive tau oligomers were more frequently observed in MCI and AD patients compared to CN subjects. Quantitative analyses indicated a decline in RGC integrity, with 46-55% and 55-56% reductions of RBPMS^+^ RGCs (P<0.01) and Nissl^+^ GCL neurons (P<0.01-0.001), respectively, in MCI and AD patients. This decrease in RGC count was accompanied by increases in necroptotic-like morphology and the cleaved caspase-3 apoptotic marker in RGCs of AD patients. Furthermore, there was a 2.1 to 3.1-fold increase (P<0.05-0.0001) in pS396-tau-laden RGCs in MCI and AD patients, with a greater abundance observed in individuals with higher Braak stages (V-VI), more severe clinical dementia ratings (CDR=3), and lower mini-mental state examination (MMSE) scores. Strong correlations were noted between the decline in RGCs and the total amount of retinal pS396-tau and pS396-tau^+^ RGCs, with pS396-tau^+^ RGC counts correlating significantly with brain neurofibrillary tangle scores (*r*= 0.71, P= 0.0001), Braak stage (*r*= 0.65, P= 0.0009), and MMSE scores (*r*= -0.76, P= 0.0004). These findings suggest that retinal tauopathy, characterized by pS396-tau and oligomeric tau in hypertrophic RGCs, is associated with and may contribute to RGC degeneration in AD. Future research should validate these findings in larger cohorts and explore noninvasive retinal imaging techniques that target tau pathology in RGCs to improve AD detection and monitor disease progression.

## Introduction

Alzheimer’s disease (AD), the most prevalent and progressive form of senile dementia, affects an estimated 6.9 million Americans aged 65 and older [1]. It is characterized by the accumulation of amyloid beta-protein (Aβ) deposits and abnormal tau protein aggregates in the brain [18, 50]. During AD progression, microtubule-associated tau proteins undergo hyperphosphorylation (p-tau) and form toxic oligomers that spread between neurons, accelerating disease progression [14, 41, 42, 54, 58, 64]. These tau species eventually aggregate into neurofibrillary tangles (NFTs) [79], disrupting cellular functions and axonal transport, which leads to synaptic dysfunction and neuronal death [83, 95, 105, 116]. The presence of abnormal tau strongly correlates with the progression of neurodegeneration and cognitive deficits in AD [20, 35, 41, 51, 65]. AD neuropathology develops many years before neurobehavioral and cognitive disturbances become salient [51, 110, 111, 124], therefore early identification of AD pathological hallmarks in the central nervous system (CNS) is crucial for early intervention and disease management.

The retina, a posterior neurosensory eye tissue, is an extension of the brain and shares many structural and functional features with the brain. New studies have revealed the genetic basis for eye-brain connections, suggesting bidirectional genetic causal links between retinal structures and neurological disorders, including AD [33, 93, 127]. Growing evidence indicates the presence of AD-related pathological features in the retinas of patients with mild cognitive impairment (MCI due to AD) and/or AD dementia, including various abnormal Aβ and tau species, vascular damage, micro- and macro-gliosis, and neurodegeneration [2, 4, 6, 7, 16, 17, 22, 27, 29–31, 37, 40, 43, 46, 48, 60–62, 67, 70, 71, 81, 85, 86, 94, 103, 104, 106–108, 113, 122, 123]. Regarding tauopathy, a wide range of abnormal tau isoforms have been identified in the retinas of AD patients, including pretangles and mature tangle forms: 3- and 4-repeat tau, p-tau and citrullinated tau forms, oligomeric tau, paired helical filaments (PHF) of tau, as well as paperclip folding of tau and NFT-like structures [27, 29, 40, 45, 46, 60, 86, 108, 122]. We recently found that the retinas of patients with MCI (due to AD) and/or AD dementia exhibit significant increases in pathogenic p-tau at specific epitopes, including S202/T205, S214, S396, S404, and T231, as well as citrullinated R209-tau and tau oligomers (T22-positive), alongside PHF^+^ and MC-1^+^ pretangle and mature tau tangles. Epitopes S199 and T212/S214 did not show such changes [108]. Moreover, oligomeric tau and pS396-tau, commonly elevated in AD brains [96, 120], were consistently increased in AD retinas and strongly associated with more severe brain pathology, advanced disease stages, and cognitive decline [108]. However, the impact of AD-related tauopathy on specific retinal cell types in patients has not yet been described.

Retinal ganglion cells (RGCs) are neurons located in the retinal ganglion cell layer (GCL; as seen in optical coherence tomography – OCT imaging) and existing in various subtypes such as midget, parasol, bistratified, and melanopsin-containing intrinsically photosensitive RGCs (mRGCs). These cells serve diverse functions, including high spatial frequency resolution, color differentiation, low spatial frequency contrast, and photoentrainment of the hypothalamus, which governs circadian rhythms [102, 121]. Dendritic protrusions from the RGC soma receive synaptic input from the axons of bipolar and amacrine cells in the inner plexiform layer (IPL). The RGCs project their axons to form the nerve fiber layer (NFL), which collects at the optic discs and continues as the optic nerve. This pathway ultimately transmits all visual information to the brain [55]. Notably, the RGCs, located in the inner retinal surface, are uniquely positioned as neurons in the CNS that can be noninvasively imaged and quantitatively assessed in vivo with high resolution using the advanced adaptive optics (AO)-OCT technology, as demonstrated in recent studies [44, 76]. This advanced imaging capability enables detailed examination of RGC pathology and may facilitate future AD diagnosis and monitoring.

In the context of AD, pioneering studies have demonstrated the loss of RGCs in patients [15, 16, 48]. Other reports have shown visual dysfunctions such as impaired contrast sensitivity, abnormal color discrimination, and diminished visual fields, which can be attributed to RGC degeneration [37, 53, 97, 100, 118]. Subsequent investigations into the AD retina found NFL thinning, reduced density of melanopsin-containing RGCs, GCL cell loss, and elevated apoptotic markers, along with intraneuronal Aβ oligomers and other Aβ species within RGCs in these patients [4–6, 23, 25, 37, 56, 60, 61, 66, 67, 70, 74]. A recent report in several transgenic murine models of AD showed RGC susceptibility, manifested as RGC dendritic field reduction, occurring in parallel with hippocampal dendritic spine loss [13]. An additional study detected an increased total tau burden in RGCs in an AD-murine model [24]. However, the vulnerability of RGCs to pathogenic tau accumulation in AD patients, particularly in the earliest stages of functional impairment (MCI due to AD), and the potential relationships with disease status, have not yet been studied.

In the current study, we addressed these gaps by first investigating the density, size, and distribution of RGCs in the superior temporal postmortem retinas of patients with MCI (due to AD) and AD dementia, compared with cognitively normal (CN) individuals. We then explored whether AD-specific pathological tau forms, pS396-tau and oligomeric tau, are present specifically in RGCs, and quantified pS396-tau-containing RGCs in this cohort. The interplay between pS396-tau-containing RGCs, retinal Aβ and tau pathology, and RGC integrity was assessed, and correlations to disease status were determined. Our analyses indicated an early and substantial decrease in RGCs, concomitant with an increase in pS396-tau-laden RGCs in MCI and AD patients compared to age- and sex-matched CN controls. The levels of total retinal pS396-tau and pS396-tau-laden RGCs correlated with the extent of RGC decline. RGCs in AD patients exhibited hypertrophic soma and nucleus displacement. Notably, increased pS396-tau^+^ RGC counts strongly correlated with corresponding brain pathology and cognitive status.

## Materials and Methods

### Postmortem Eyes

Human eye and brain tissues were collected from donor patients with premortem clinical diagnoses of MCI and AD dementia (confirmed by postmortem AD neuropathology), along with age- and sex-matched CN controls (total *n*=41 subjects). These tissues were primarily obtained from the Alzheimer’s Disease Research Center (ADRC) Neuropathology Core in the Department of Pathology (IRB protocol HS-042071) at the Keck School of Medicine, University of Southern California (USC, Los Angeles, CA). Additional eyes were obtained from the National Disease Research Interchange (NDRI, Philadelphia, PA) under the approved Cedars-Sinai Medical Center IRB protocol Pro00019393. Both USC-ADRC and NDRI maintain human tissue collection protocols approved by their managerial committees and subject to oversight by the National Institutes of Health. Histological studies at Cedars-Sinai Medical Center were performed under IRB protocols Pro00053412 and Pro00019393. Demographic, clinical, and neuropathological information on human donors is detailed in **Table 1 and Suppl. Table 1**. Subjects with macular degeneration, glaucoma, and diabetic retinopathy were excluded from this study. The available retinal tissues from individual donors are specified in **Suppl. Table 1**. For the histopathological analysis, the human cohort consisted of AD dementia (n=15), MCI due to AD (n=10), and CN controls (n=16). All patients’ identities were protected by de-identifying tissue samples, ensuring they could not be traced back to the donors.

**Table 1.**
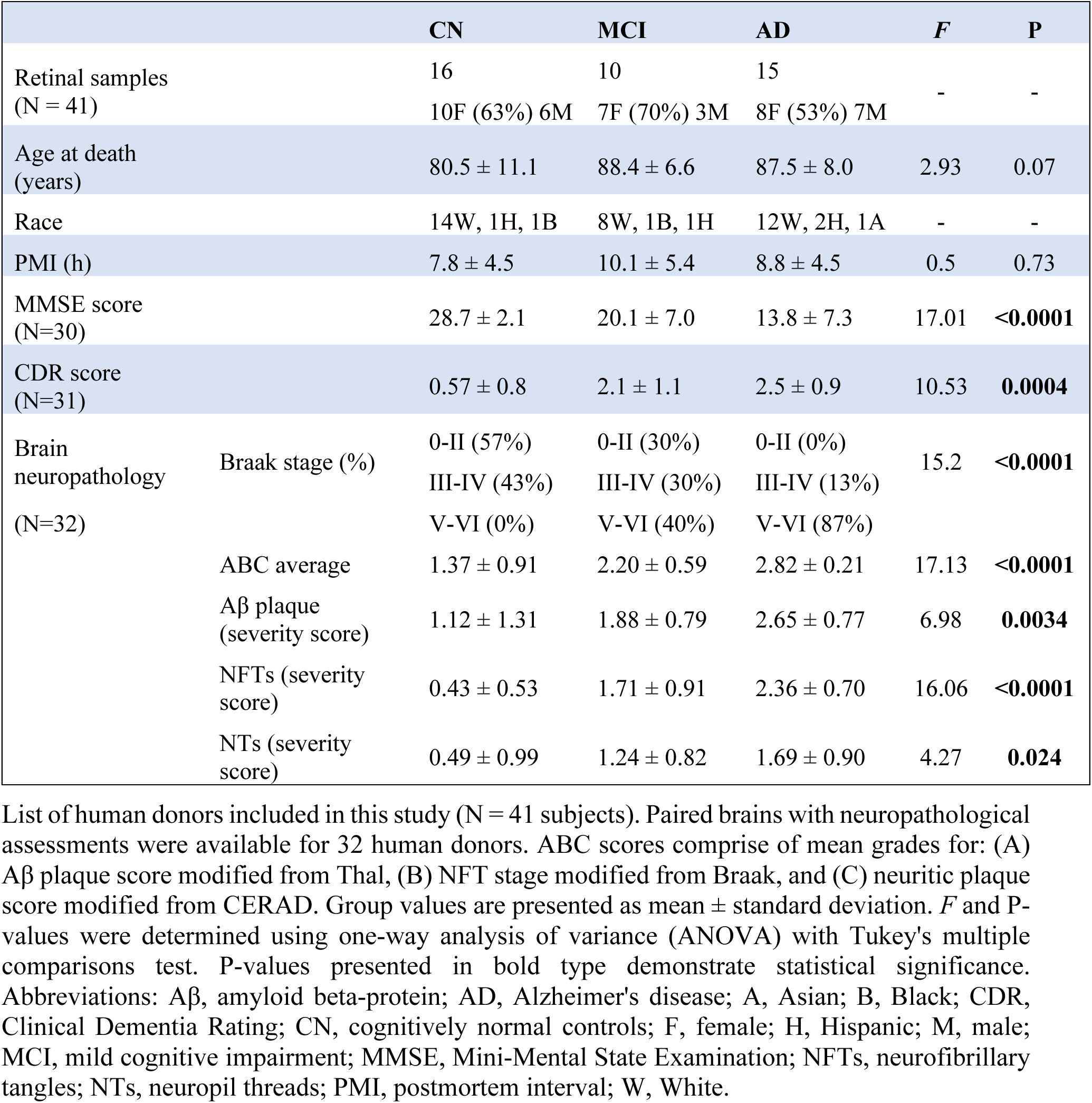
Demographic and neuropathological data on human brain and retinal donors in this study.

|  |  | CN | MCI | AD | *F* | *P* |
| --- | --- | --- | --- | --- | --- | --- |
| Retinal samples<br>(N = 41) |  | 16 | 10 | 15 | - | - |
|  |  | 10F (63%) 6M | 7F (70%) 3M | 8F (53%) 7M |  |  |
| Age at death<br>(years) |  | 80.5 ± 11.1 | 88.4 ± 6.6 | 87.5 ± 8.0 | 2.93 | 0.07 |
| Race |  | 14W, 1H, 1B | 8W, 1B, 1H | 12W, 2H, 1A | - | - |
| PMI (h) |  | 7.8 ± 4.5 | 10.1 ± 5.4 | 8.8 ± 4.5 | 0.5 | 0.73 |
| MMSE score<br>(N=30) |  | 28.7 ± 2.1 | 20.1 ± 7.0 | 13.8 ± 7.3 | 17.01 | **<0.0001** |
| CDR score<br>(N=31) |  | 0.57 ± 0.8 | 2.1 ± 1.1 | 2.5 ± 0.9 | 10.53 | **0.0004** |
| Brain<br>neuropathology<br>(N=32) | Braak stage (%) | 0-II (57%) | 0-II (30%) | 0-II (0%) | 15.2 | **<0.0001** |
|  |  | III-IV (43%) | III-IV (30%) | III-IV (13%) |  |  |
|  |  | V-VI (0%) | V-VI (40%) | V-VI (87%) |  |  |
|  | ABC average | 1.37 ± 0.91 | 2.20 ± 0.59 | 2.82 ± 0.21 | 17.13 | **<0.0001** |
|  | Aβ plaque<br>(severity score) | 1.12 ± 1.31 | 1.88 ± 0.79 | 2.65 ± 0.77 | 6.98 | **0.0034** |
|  | NFTs (severity<br>score) | 0.43 ± 0.53 | 1.71 ± 0.91 | 2.36 ± 0.70 | 16.06 | **<0.0001** |
|  | NTs (severity<br>score) | 0.49 ± 0.99 | 1.24 ± 0.82 | 1.69 ± 0.90 | 4.27 | **0.024** |

List of human donors included in this study (N = 41 subjects). Paired brains with neuropathological assessments were available for 32 human donors. ABC scores comprise of mean grades for: (A) Aβ plaque score modified from Thal, (B) NFT stage modified from Braak, and (C) neuritic plaque score modified from CERAD. Group values are presented as mean ± standard deviation. *F* and *P*-values were determined using one-way analysis of variance (ANOVA) with Tukey's multiple comparisons test. *P*-values presented in bold type demonstrate statistical significance. Abbreviations: Aβ, amyloid beta-protein; AD, Alzheimer's disease; A, Asian; B, Black; CDR, Clinical Dementia Rating; CN, cognitively normal controls; F, female; H, Hispanic; M, male; MCI, mild cognitive impairment; MMSE, Mini-Mental State Examination; NFTs, neurofibrillary tangles; NTs, neuropil threads; PMI, postmortem interval; W, White.

### Clinical and Neuropathological Assessments

The ADRC provided clinical and neuropathological reports on patients’ neurological examinations, neuropsychological and cognitive tests, family history, and medication lists, as collected in the ADRC system using the Uniform Data Set (UDS) [12]. The NDRI provided the medical history of additional patients. Most cognitive evaluations were performed annually and, in most cases, less than one year prior to death. Cognitive testing scores from evaluations made closest to the patient’s death were used for this analysis. Two global indicators of cognitive status were used for clinical assessment: the Clinical Dementia Rating (CDR scores: 0 = normal; 0.5 = very mild impairment; 1 = mild dementia; 2 = moderate dementia; or 3 = severe dementia) [82] and the Mini-Mental State Examination (MMSE scores: 24–30 = CN; 20–23 = MCI; 10–19 = moderate dementia; or 9 ≥ severe dementia) [34]. In this study, the composition of the clinical diagnostic groups (AD, MCI, or CN) was determined by source clinicians based on a comprehensive battery of tests, including neurological examinations, neuropsychological evaluations, and the cognitive tests. Specifically, the diagnosis of MCI due to AD was assigned to patients who had an antemortem clinical diagnosis of MCI (based on the comprehensive battery of behavioral and cognitive tests) caused by AD. These patients had a postmortem confirmation of AD neuropathology (according to the ADNC—Alzheimer’s disease neuropathological change guidelines) and showed no evidence of other diseases, such as Lewy body dementia, Parkinson’s disease, FTD/FTLD (PSP or Pick’s disease), or cognitive impairment due to stroke or small vessel disease.

To obtain a final diagnosis based on the neuropathological reports, we used the modified Consortium to Establish a Registry for Alzheimer’s Disease (CERAD) criteria [77, 99], as outlined in the National Institute on Aging (NIA)/Regan protocols with revisions by the NIA and Alzheimer’s Association [49]. The assessment included Aβ burden (measured as diffuse, immature, or mature plaques), amyloid angiopathy, neuritic plaques, NFTs, neuropil threads (NTs), granulovacuolar degeneration, Lewy bodies, Hirano bodies, Pick bodies, balloon cells, neuronal loss, microvascular changes, and gliosis. These pathologies were assessed in multiple brain areas, including the hippocampus (particularly the Cornu ammonis CA1, at the level of the thalamic lateral geniculate body), entorhinal cortex, superior frontal gyrus of the frontal lobe, superior temporal gyrus of the temporal lobe, superior parietal lobule of the parietal lobe, primary visual cortex (Brodmann Area-17), and visual association (Area-18) of the occipital lobe. In all cases, uniform brain sampling was conducted by a neuropathologist.

Cerebral amyloid plaques, NFTs, and NTs were evaluated using anti–β-amyloid mAb clone 4G8 immunostaining, Thioflavin-S (ThioS) histochemical staining, and Gallyas silver staining in formalin-fixed, paraffin-embedded tissue sections. The ADRC neuropathologists assigned severity scores based on semi-quantitative observations. The scale for Aβ/neuritic plaques was determined by the presence of 4G8- and/or Thioflavin-S-positive and/or Gallyas silver-positive plaques measured per 1 mm^2^ of brain area (0 = none; 1 = sparse [≤ 5 plaques]; 3 = moderate [6–20 plaques]; 5 = abundant/frequent [21–30 plaques or greater]; or N/A = not applicable), as previously described [80] in the NACC NP Guidebook, Version 10, January 2014: https://naccdata.org/data-collection/forms-documentation/np-10. The brain NFT or NT severity scoring system was derived from observed burden of these AD neuropathologic changes, as detected by Gallyas silver and/or Thioflavin-S staining [79, 80, 119], and measured per 1 mm^2^ of brain area. The assigned NFT or NT scores were as follows: 0 = none; 1 = sparse (mild burden); 3 = moderate (intermediate burden); or 5 = frequent (severe burden). For both histochemical and immunohistochemical staining, each anatomical area of interest was assessed for relevant pathology using a 20X objective (200X high power magnification), and representative fields were graded using the semiquantitative scale as detailed above. Validation of AD neuropathic change (ADNC), especially NTs, was performed using a 40X objective (400X high power magnification), and an average of two readings was assigned to each individual patient.

A final diagnosis of AD neuropathological change was determined using an “ABC” score derived from three separate 4-point scales. We used the modified Aβ plaque Thal score (A0 = no Aβ or amyloid plaques; A1 = Thal phase 1 or 2; A2 = Thal phase 3; or A3 = Thal phase 4 or 5) [115]. For the NFT stage, we applied the modified Braak staging for silver-based histochemistry or p-tau IHC (B0 = no NFTs; B1 = Braak stage I or II; B2 = Braak stage III or IV; or B3 = Braak stage V or VI) [19]. For neuritic plaques, we used the modified CERAD score (C0 = no neuritic plaques; C1 = CERAD score sparse; C2 = CERAD score moderate; or C3 = CERAD score frequent) [77]. Neuronal loss, gliosis, granulovacuolar degeneration, Hirano bodies, Lewy bodies, Pick bodies, and balloon cells were all evaluated (0 = absent; 1 = present) in multiple brain areas by staining tissues with hematoxylin and eosin (H&E). Brain atrophy was evaluated (0 = none; 1 = mild; 3 = moderate; 5 = severe; or 9 = not applicable).

### Processing of Eye Globes and Retinal Tissues

The processing of eye globes, isolation and preparation of retinal strips, and retinal immunostaining were extensively detailed in [60, 61, 108]. Briefly, donor eyes were collected within an average of 9 hours after death and subjected to the following preservation methods: 1) preserved in Optisol-GS media (Bausch & Lomb, 50006-OPT) and stored at 4°C for less than 24 hours, or 2) punctured once and fixed in 10% neutral buffered formalin (NBF) or 4% paraformaldehyde (PFA) and stored at 4°C. Regardless of the source of the human donor eye (USC-ADRC or NDRI), the same tissue collection and processing methods were applied.

### Preparation of Retinal Strips

Eyes fixed in 10% NBF or 4% PFA were dissected as previously described [60, 61, 108]. Flatmounts were prepared after careful dissection of the eye globes and thorough cleaning of the vitreous humor. Flatmount strips (∼2 mm wide) extending diagonally from the optic disc (OD) to the ora serrata (∼20–25 mm long) were prepared in 4 predefined regions: Superior Temporal (ST), Inferior Temporal (IT), Inferior Nasal (IN), and Superior Nasal (SN). In this study, we focused our analysis on the ST retinal strip due to the high presence of AD pathology in this region [60, 61, 108]. The flatmount-derived strips were then paraffinized using standard techniques and embedded in paraffin after flip-rotating 90° horizontally. The retinal strips were sectioned (7-10 µm thick) and mounted on microscope slides coated with APES. This sample preparation technique allowed for extensive and consistent access to retinal quadrants, layers, and pathological subregions.

### Immunofluorescent Staining

Retinal sections were deparaffinized using 100% xylene twice (10 minutes each), rehydrated with decreasing concentrations of ethanol (100% to 70%), and washed with distilled water followed by PBS. After deparaffinization, tissue sections were treated with target retrieval solution (pH 6.1; S1699, Dako) at 98°C for 1 hour and then washed with PBS. Next, tissues were incubated in blocking buffer (Dako #X0909) supplemented with 0.1% Triton X-100 (Sigma, T8787) for 1 hour at room temperature (RT), followed by overnight incubation with primary antibody (Ab) at 4°C (Abs information provided in **Suppl. Table 2**). The sections were then washed three times with PBS and incubated with secondary Abs against each species (1:200, **Suppl. Table 2**) for 1 hour at RT. After rinsing with PBS three times, the sections were mounted with ProLong Gold antifade reagent with DAPI (Thermo Fisher #P36935).

### Peroxidase-based Immunostaining

After deparaffinization and antigen retrieval treatment, the tissues were treated with 70% formic acid (ACROS) for 10 minutes at room temperature. The tissues were then washed with wash buffer (Dako S3006) supplemented with 0.1% Triton X-100 (Sigma, T8787) for 1 hour, followed by treatment with H_2_O_2_ for 10 minutes and a rinse with wash buffer. Primary Ab (**Suppl. Table 2**) were diluted with background reducing components (Dako S3022) and incubated with the tissues overnight at 4°C. The tissues were rinsed thrice with wash buffer on a shaker and incubated for 30 minutes at 37°C with secondary Ab (goat anti-rabbit HRP conjugated, Dako Envision K4003), followed by three more rinses with wash buffer on a shaker. Diaminobenzidine (DAB) substrate (Dako K3468) was then applied. Some slides were counterstained with hematoxylin and mounted with Faramount aqueous mounting medium (Dako, S3025). Routine controls were processed using an identical protocol, while omitting the primary antibodies to assess nonspecific labeling.

### Nissl Staining

A basic (alkaline) dye was used to label nuclei and granules (i.e., ribosomal RNA) in neurons. The cytoplasm of neurons is specifically stained with the Nissl staining technique, while the <u>perikarya</u> of other cellular elements are either weakly visualized or not at all [52]. Deparaffinized and rehydrated sections were stained in 0.1% Cresyl Violet acetate (Sigma #C5042) for 5 min, rapidly rinsed in tap water, and briefly dipped in 70% ethanol. The sections were then dehydrated through 2 changes of absolute ethanol for 3 minutes each, followed by immersion in xylene twice for 2 minutes and mounted in mounting medium xylene (Fisher scientific company, L.L.C. #245-691). An average of 12 images (from the superior quadrant), covering the retinal neurons from the optic disc to the ora serrata, were captured at a 20x objective and analyzed to quantify the area and number of retinal GCL neurons.

### Microscopy and Stereological Quantification

Fluorescence and brightfield images were acquired using a Carl Zeiss Axio Imager Z1 fluorescence microscope (with motorized Z-drive) equipped with ApoTome, AxioCam HRc, and AxioCam MRm monochrome cameras (version 3.0; resolution of 1388 × 1040 pixels, 6.45 µm × 6.45 µm pixel size, and a dynamic range of >1:2200, which delivers low-noise images due to a Peltier-cooled sensor) with ZEN 2.6 blue edition software (Carl Zeiss MicroImaging, Inc.). Multi-channel image acquisition was used to create images with multiple channels. Images were consistently captured at the same focal planes with identical exposure time, using a 20x objective at a resolution of 0.25 µm. Approximately 15 images were obtained from each retina. The acquired images were converted to grayscale and standardized to baseline using a histogram-based threshold in Fiji ImageJ (NIH) software (version 1.53c). For each biomarker, the total area of immunoreactivity was determined using the same threshold percentage from the baseline in ImageJ (with the same percentage threshold setting for all diagnostic groups). The images were then subjected to particle analysis to determine the immunoreactive (IR) area and/or area fraction (%).

### RGC Soma Size Measurement

The size of RGC somas was measured using Fiji ImageJ (NIH) software (version 1.53c) with the polygonal selection tool. For each 20x retinal image, the soma area of up to three cells was manually assessed, focusing on the three largest cells in each field. The average soma area for each subject was then computed, followed by statistical analysis. On average, 30 somas were analyzed per patient, with a total of 542 somas measured.

### Statistical Analysis

GraphPad Prism Software version 9.5.1 was used for statistical analyses. One-way or two-way ANOVA followed by Tukey’s multiple comparison post-test was used to determine statistical significance between three or more groups. Two-group comparisons were analyzed using a two-tailed unpaired Student’s *t-test*. The statistical association between two or more Gaussian-distributed variables was determined by Pearson’s correlation coefficient (*r*) test. Scatterplot graphs present the null hypothesis of pair-wise Pearson’s *r*, with unadjusted P values indicating the direction and strength of the linear relationship between two variables. Results are expressed as the mean ± standard deviation (SD) in tables and as median, lower, and upper quartiles in violin plots. Degrees of significance are presented as: *P < 0.05, **P < 0.01, ***P < 0.001, and ****P < 0.0001. Data analysis was conducted using coded identifiers, and analysts remained blinded to the diagnostic groups until all analyses were completed.

## Results

To investigate the integrity of RGCs, including their number, morphology, and distribution in relation to abnormal retinal tau isoforms and their accumulation within RGCs in early and advanced-stage AD, we selected and analyzed retinal superior temporal (ST) cross-sections (**Fig. 1a, b**) from patients with MCI due to AD (n=10, mean age 88.4 ± 6.6 years, 7 females/3 males) and AD dementia (n=15, mean age 87.5 ± 8.0 years, 8 females/7 males), compared to CN controls (n=16, mean age 80.5 ± 11.1 years, 10 females/6 males). Demographic, clinical, and neuropathological information are detailed in **Table 1** (list of individual donor eyes and respective brains detailed in **Suppl. Table 1**).

**Figure 1.**
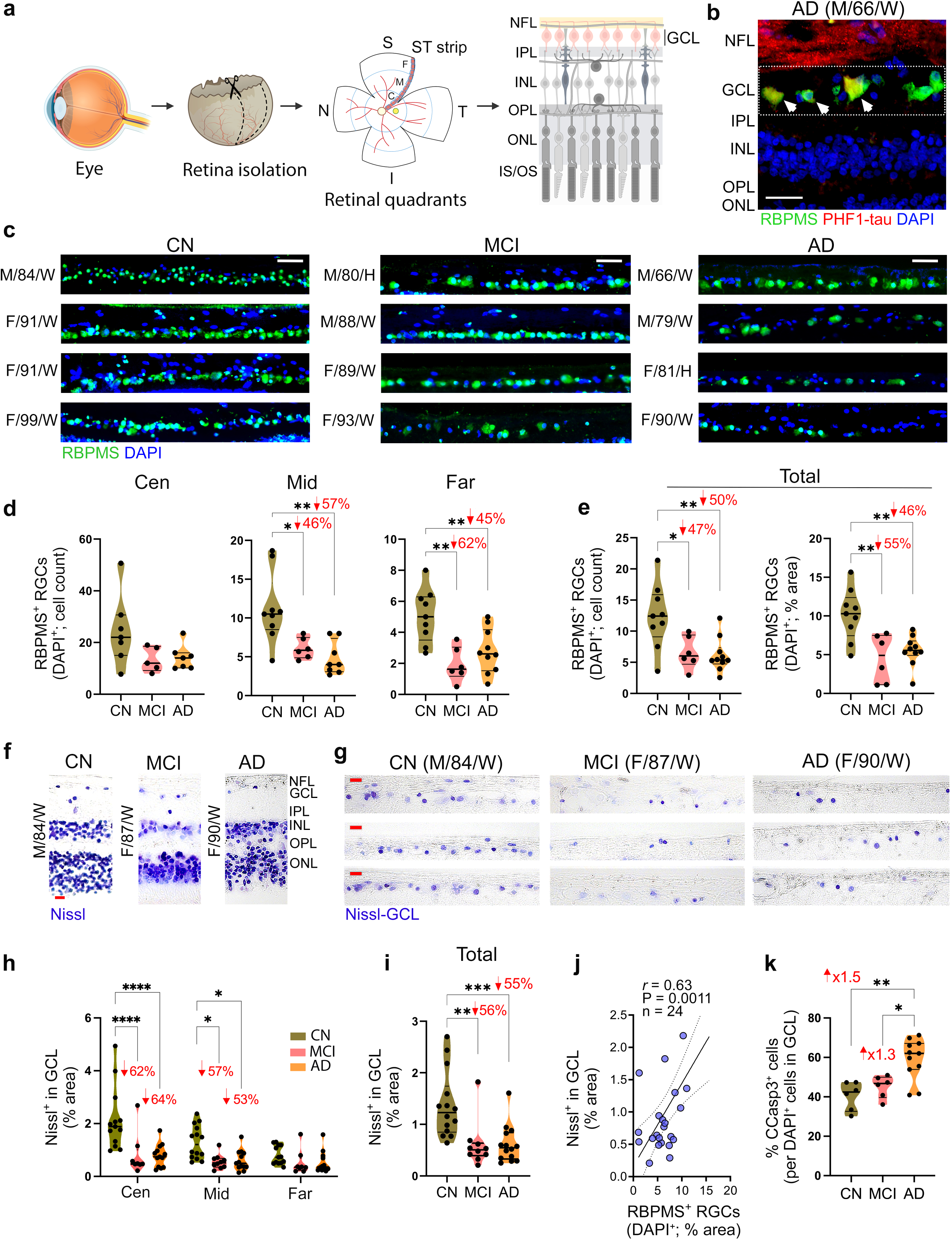
Ganglion cell integrity in retinal tissues of MCI and AD patients. **a** Illustration of the histological process, including retinal isolation, cross-section preparation, and analysis of the superior temporal (ST) strip, extending from the optic disc to the ora serrata and anatomically predefined into central (Cen), middle (Mid) and far-peripheral (Far) subregions. The retinal ganglion cell layer (GCL) was analyzed in this study. **b** Microscopic image of a retinal cross-section from an AD patient, immunolabeled with retinal ganglion cell (RGC)-specific marker, ribonucleic acid binding protein with multiple splicing (RBPMS; green), and paired-helical filament of tau (PHF1-tau; red), along with nuclei labelling with DAPI (blue). Scale bar: 25 µm. **c** Representative microscopic images of RBPMS^+^ RGCs within the GCL, labeled with RBPMS (green) and DAPI (blue), in retinal cross-sections from patients with mild cognitive impairment (MCI due to AD, n=4) and Alzheimer’s disease (AD) dementia (n=4), and cognitively normal individuals (CN, n=4). Scale bar: 50 µm. **d, e** Violin graphs display the quantitative immunohistochemistry analyses of RBPMS^+^DAPI^+^ RGCs by **d** cell count in Cen, Mid- and Far-peripheral subregions, and **e** cell count (left) and percent area (right) in the total ST region (n=25 subjects; n=9 CN, n=6 MCI, n=10 AD). **f, g** Representative microscopic images of retinal cross-sections from CN, MCI, and AD donors labeled with Nissl stain (purple) in **f** all analyzed retinal layers (ONL to NFL) and **g** GCL separately. Scale bars: 20 µm. **g, h** Quantitative analyses of Nissl^+^ percent area in GCL in **g** the Cen, Mid, and Far-peripheral subregions (n=33-37) and **h** the total ST region (n=38 subjects; n=14 CN, n=10 MCI, n=14 AD). **i** Pearson’s correlation coefficient (*r*) analysis between RBPMS^+^ RGCs percent area and Nissl^+^ cells (in GCL) percent area. **j** Quantitative analysis of the percent area of early apoptotic cell marker, cleaved caspase-3 (CCasp3)^+^ in GCL, normalized to nuclei count (n=23 subjects; n=6 CN, n=6 MCI, n=11 AD). Individual data points and median, lower and upper quartiles are shown in violin plots. *P < 0.05, **P < 0.01, ***P < 0.001, ****P < 0.0001, by one-way or two-way ANOVA followed by Tukey’s post-hoc multiple comparison test. Percent decreases and fold changes are shown in red. F, female; M, male; Age (in years); Ethnicity: W, White and H, Hispanic; NFL, Nerve fiber layer; IPL, Inner Plexiform Layer; INL, Inner Nuclear Layer; OPL, Outer Plexiform Layer; ONL, Outer Nuclear Layer; IS/OS, inner segment and outer segment. Illustrations created with Biorender.com.

### 1. Severe RGC decline in MCI and AD patients

We first assessed RGC numbers and distribution across ST subregions in a sub-cohort of patients with MCI (n=6, mean age 89.5 ± 5.24 years, 3 females/3 males), AD (n=10, mean age 86.0 ± 8.89 years, 4 females/6 males), and age- and sex-matched CN controls (n=9, mean age 85.89 ± 11.85 years, 5 females/4 males), using a selective pan-RGC marker, ribonucleic acid binding protein with multiple splicing (RBPMS), for immunohistochemical (IHC) analysis. According to previous studies, RBPMS is specifically expressed in the entire RGC population, despite the heterogeneity of other neurons under pathological conditions, including displaced amacrine cells within the GCL [88, 91, 98]. In comparison to the retinas of CN individuals, the density of RGCs was lower in MCI and AD dementia patients, with their cytoplasm appearing enlarged or swollen (**Fig. 1c** and **Suppl. Figure 1a**). Analysis of RBPMS^+^ RGC cell counts per retinal subregion (central, mid-, and far-periphery) and the total ST region revealed substantial reductions in RGC count and percent area–ranging from 42% to 65%–in MCI and AD patients compared to CN individuals (**Fig. 1d, e** and **Suppl. Fig. 1b**; P<0.05-0.01). RGC loss in MCI and AD retinas appeared more extensive in the mid- and far-periphery regions, which are further distal from the optic nerve head.

We next examined neurodegeneration in the GCL using histological Nissl staining, an alkaline dye that labels nuclei and granules (i.e., ribosomal RNA) in neurons, in a sub-cohort of patients diagnosed with MCI (n=10, mean age 88.4 ± 6.6 years, 7 females/3 males), AD (n=15, mean age 87.5 ± 8.0 years, 8 females/7 males), and CN controls (n=14, mean age 80.6 ± 12.1 years, 9 females/5 males) (**Fig. 1f-i**). Representative images showed a reduction in cells numbers across all retinal layers (**Fig. 1f**), particularly in the GCL (**Fig. 1g**), in MCI and AD patients compared to CN controls. Quantitative analysis of Nissl^+^ percent area in the GCL across retinal subregions indicated a marked 53%-64% neuronal loss in the central and mid-peripheral subregions of MCI and AD patients compared to CN controls (**Fig. 1h**); no statistically significant reduction was observed in the far-peripheral subregion. In the total ST region, a substantial 55%-56% reduction in Nissl^+^ percent area in the GCL was observed for both AD and MCI groups compared to CN controls (**Fig. 1i**; P<0.01-0.001). Pearson’s correlation coefficient (*r*) analysis demonstrated a strong correlation between the two RGC integrity parameters, RBPMS^+^ RGCs and Nissl^+^ neurons in the GCL (*r*=0.63, P=0.0011; **Fig. 1j**). To assess whether GCL residing neurons were lost due to apoptotic cell death mechanisms, we performed IHC using an antibody against cleaved caspase 3 (CCasp3), an early apoptotic marker [117]. Analysis of the percent of CCasp3^+^ cells in the GCL revealed a significant 1.5-fold and 1.3-fold increase in AD retinas compared to CN and MCI retinas, respectively (**Fig. 1k**; P<0.05-0.01), with no differences noted between the MCI and CN retinas. When the CCasp3^+^ cell immunoreactive area in the total retina was normalized to retinal thickness, there were highly significant 3.1-fold and 2-fold increases in AD compared to CN and MCI, respectively (P<0.001-0.0001), with a trend toward a 1.5-fold increase in MCI compared to CN, reaching significance by Student’s *t*-test (**Suppl. Fig. 1c**).

### 2. Increased pS396-tau laden RGCs of MCI and AD patients is linked to RGC hypertrophy and loss

We recently found significant increases in AD-related tau isoforms, particularly pretangles such as pS396-tau and tau oligomers (oligo-tau), in the retinas of MCI and AD patients, which strongly correlated with corresponding brain pathology and cognitive deficits [108]. In this study, we investigated whether RGCs are vulnerable to these tau isoforms in early and advanced AD (**Fig. 2**; extended data in **Suppl. Fig. 2**). Utilizing the same sub-cohort of patients outlined above for the RBPMS analysis, we performed an IHC analysis employing a combination of RBPMS and pS396-tau, which recognizes the hyperphosphorylated tau protein at serine residue 396 in the C-terminal region. Representative microscopic images depicted increases in pS396-tau burden within the OPL, IPL, GCL, and NFL, along with cell swelling (hypertrophic soma), whereas a reduction in the number of RBPMS^+^ RGC was observed in MCI and AD patients was seen compared to CN controls (**Fig. 2a)**. The three-parallel-string staining pattern of retinal pS396-tau in the IPL of MCI and AD patients appeared to accumulate in neuronal dendrites of RGCs connecting with axons of bipolar and amacrine cells. Notably, morphological changes were observed in the RGCs of MCI and AD patients compared to CN controls (**Fig. 2a-d** and **Suppl. Fig. 2a, b**). These ganglion cells exhibited granulovacuolar vesicles degeneration (GVD)-like bodies and nucleus displacement, as indicated by white and red arrows, respectively (**Fig. 2b** and **Suppl. Fig. 2a**). Analysis of the enlarged and granulomatous soma areas of RBPMS^+^ RGCs revealed a significant 1.5-fold increase in RGC soma size in AD patients compared to CN controls (P=0.018), with no difference observed in RGC size in MCI patients (**Fig. 2b’**).

**Figure 2.**
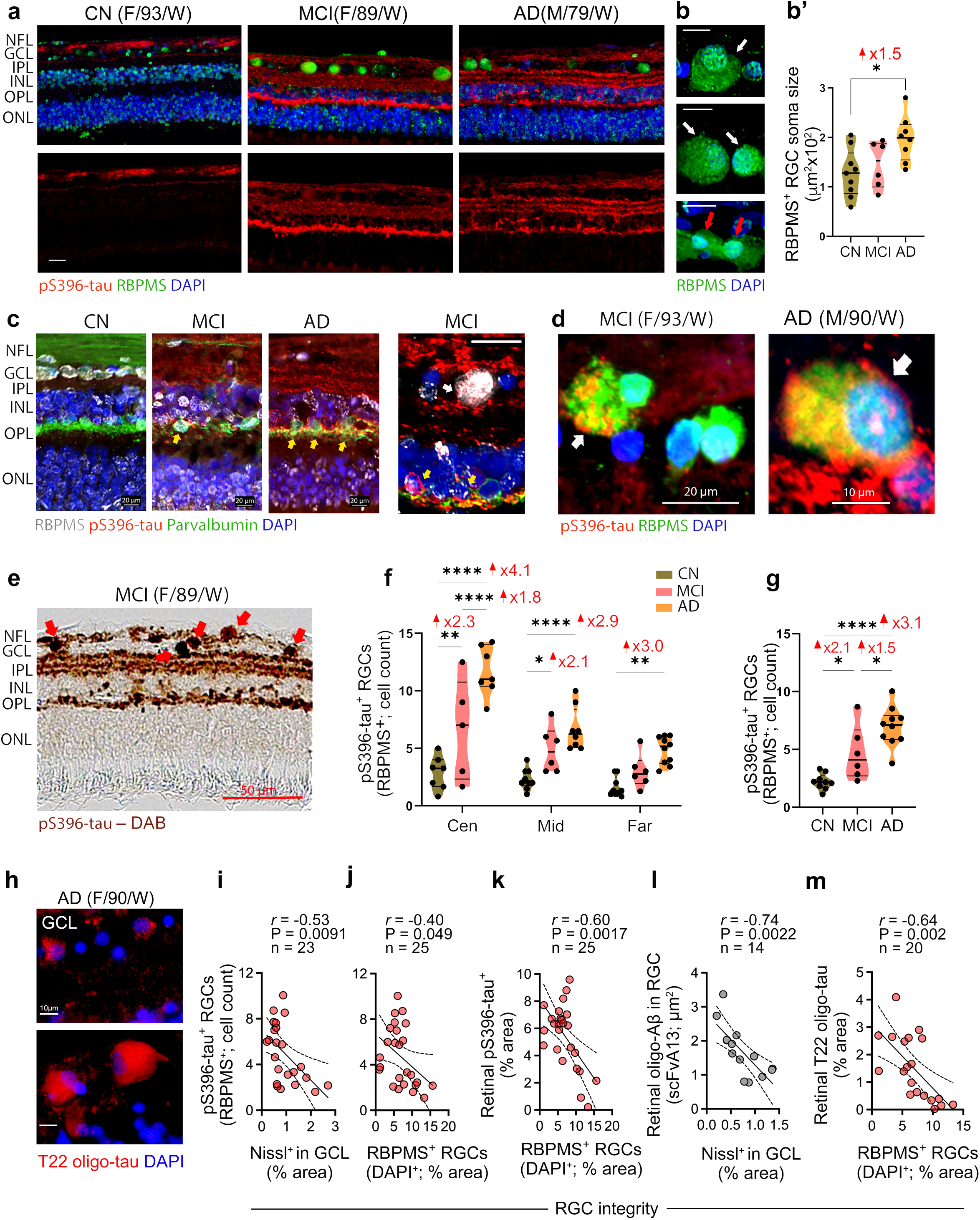
Pretangle tau pathology in RGCs of MCI and AD patients. **a** Representative microscopic images of retinal cross-sections immunofluorescently stained for hyperphosphorylated (p)tau at S396 epitope (pS396-tau, red), RGC-specific marker, RBPMS (green) and nuclei (DAPI, blue) in retinal cross-sections from patients with mild cognitive impairment (MCI due to AD) and Alzheimer’s disease (AD) dementia, and cognitively normal individuals (CN). **a, b** Retinas from MCI and AD patients exhibit increases in pS396-tau isoforms and the RGCs exhibit reduced numbers, a hypertrophic cytoplasm (cell soma swelling), and abnormal morphology, including granulovacuolar vesicles degeneration (GVD)-like bodies and nucleus displacement (white arrows indicate enlarged and granulomatous soma area and red arrows point to nuclear displacement). Scale bars: 20µm. **b’** Quantitative analysis of RBPMS^+^ RGC soma cell size in patients with MCI (n=6) and AD (n=8), and in CN controls (n=9). **c** Representative immunofluorescent images of retinal cross-section labelled for RBPMS RGCs (white), pS396-tau (red), amacrine and RGCs marker - parvalbumin (green), and nuclei (DAPI, blue) in CN, MCI and AD subjects. Colocalization of pS396-tau in parvalbumin^+^ amacrine cells (yellow arrows) and RBPMS^+^ RGCs (white arrows) are shown. Scale bars: 20µm. **d** High-magnification microscopic images depicting pS396-tau accumulation (red) in swollen RBPMS^+^ RGCs (green) with hypertrophic soma (white arrows). Scale bar sizes are indicated on images. **e** Representative microscopic image of peroxidase-based staining for pS396-tau isoforms (brown) within retinal layers, and specifically, in RGCs of a MCI patient. Scale bar: 50 µm. **f, g** Cell count of pS396-tau^+^ RGCs in retinal **f** Cen, Mid- and Far-peripheral subregions (n=19-25) and **g** total ST region (n=9 CN, n=6 MCI, n=10 AD). **h** Representative images of T22^+^ oligo-tau in the GCL of an AD patient. Scale bars: 10 µm. **i-m** Pearson’s correlation coefficient (*r*) analyses between: **i** % area of Nissl^+^ in GCL or **j** RBPMS^+^ RGCs % area and pS396-tau^+^ RGC count, **k** RBPMS^+^ RGC % area and retinal pS396-tau^+^ % area, **l** % area of Nissl^+^ in GCL and retinal scFvA13^+^Aβ (oligo-Aβ) in RGCs, and **m** RBPMS^+^ RGC % area and retinal T22^+^ tau oligomers (oligo-tau). Individual data points and median, lower and upper quartiles are shown in violin plots. *P < 0.05, **P < 0.01, ****P < 0.0001, by one-way or two-way ANOVA followed by Tukey’s post-hoc multiple comparison test. Fold changes are shown in red. F, female; M, male; Age (in years); Ethnicity: W, White; NFL, Nerve fiber layer, GCL, ganglion cell layer; IPL, Inner Plexiform Layer; INL, Inner Nuclear Layer, OPL, Outer Plexiform Layer; ONL, Outer Nuclear Layer; RGC, Retinal ganglion cells.

To assess whether p-tau inclusions exist and increase within RGCs of MCI and AD patients, we next immunolabeled retinal cross-sections for pS396-tau in combination with RBPMS and parvalbumin, the latter being a marker of horizontal cells within the OPL and RGCs [57]. Our analysis identified pS396-tau accumulation within hypertrophic RBPMS^+^ RGCs and horizontal cells of MCI and AD patients, and occasionally in RBPMS^+^ RGCs of CN individuals (**Fig. 2c, d** and **Suppl. Fig. 2b**). Moreover, pS396-tau build-up within the somas of RGCs in the GCL was evident in non-fluorescence, peroxidase-based IHC staining (**Fig. 2e**, red arrows). Quantitative analysis of retinal cross-sections in this cohort showed a highly significant 2.4-fold increase in total pS396-tau^+^ % area in MCI and AD patients compared to CN controls (**Suppl. Fig. 2c**; p<0.0001). Importantly, compared to the CN retina, pS396-tau-positive RBPMS^+^ RGC counts were significantly increased in MCI (2.1-2.3-fold; P<0.05-0.01), and AD (2.9-4.1-fold; P<0.01-0.0001) retinas when analyzed per retinal subregion and in the total ST region (**Fig. 2f, g**). Increases in the pS396-tau^+^ RBPMS^+^ RGC count, as well as the percentage area of pS396-tau^+^ in the GCL of MCI and AD patients, were more significant in the central ST retina (**Fig. 2f** and **Suppl. Fig. 2d, e**). Additional analysis of T22^+^ oligo-tau in the retinas of MCI and AD patients compared to CN controls identified oligo-tau aggregates within swollen RGCs of the GCL (**Fig. 2h** and **Suppl. Fig. 2f**).

To investigate the interrelations between pS396-tau-containing RBPMS^+^ RGCs, retinal pS396-tau, retinal Aβ burden, retinal oligo-tau, and RGC loss, we applied Pearson’s correlation coefficient (*r*) analyses (**Fig. 2i-m** and **Suppl. Fig. 2g-j**) in our cohort. As expected, we found a strong positive correlation between retinal pS396-tau burden and the number of pS396-tau^+^ RBPMS^+^ RGCs (**Suppl. Fig. 2g**; *r*=0.72 and P<0.0001). An unexpected strong correlation was detected between retinal Aβ_42_ burden and pS396-tau^+^ RBPMS^+^ RGC number (**Suppl. Fig. 2h**; *r*=0.69 and P=0.003). We then assessed whether there was a connection between the presence of pS396-tau in RBPMS^+^ RGCs, RGC loss, and the level of CCasp3^+^ in the GCL. Pearson’s correlation analyses revealed moderate associations between pS396-tau^+^ RBPMS^+^ RGCs and GCL Nissl^+^ cells (**Fig. 2i**; *r*=-0.53 and P=0.0091), or RBPMS^+^ RGCs (**Fig. 2j**; *r*=-0.40 and P=0.049). Notably, the apoptotic marker CCasp3^+^ cells in the GCL strongly correlated with pS396-tau^+^ RBPMS^+^ RGCs (**Suppl. Fig. 2i**; *r*=0.66 and P=0.036). We next assessed the relationship between overall retinal pS396-tau^+^, retinal Aβ_42_, retinal intra-RGC Aβ oligomers, and retinal oligo-tau burdens and RGC integrity. Pearson’s correlation analyses revealed moderate to strong correlations between retinal pS396-tau^+^ (**Fig. 2k**; *r*=-0.60 and P=0.0017), retinal 12F4^+^-Aβ_42_ (**Suppl. Fig. 2l**; *r*=-0.53 and P=0.033), retinal scFvA13^+^Aβ oligomers in RGCs (**Fig. 2l**; *r*=-0.74 and P=0.0022), and retinal T22^+^ tau oligomers (**Fig. 2m** *r*=-0.64, P=0.002), with RGC reduction.

### 3. Retinal p-tau-containing ganglion cells correlate with AD status

We further tested the potential relationship between pS396-tau^+^ RBPMS^+^ RGCs or diminished RGCs and the severity of brain pathology and cognitive deficits (**Fig. 3**, **Tables 2-3**; extended data in **Suppl. Fig. 3**). Pearson’s correlation coefficient (*r*) analyses revealed that pS396-tau^+^ RGC count strongly associated with brain Aβ-plaque and NFT severity scores (**Fig. 3a, b**; *r*=0.62, P=0.0017 and *r*=0.71, P=0.0001, respectively). Stratifying patients based on Braak stage severity showed significant 1.9-2.7-fold increases in pS396-tau^+^ RBPMS^+^ RGCs in the high (V-VI) and, to a lesser extent, the intermediate (III-IV) Braak stage groups compared to the low (0-II) group (**Fig. 3c**; P=0.0033 and P=0.042, respectively). The pS396-tau^+^ RBPMS^+^ RGCs were strongly correlated with Braak stage (**Fig. 3d**; *r*=0.65, P=0.0009), while no correlation was detected between RBPMS^+^ RGC count and Braak stage (**Suppl. Fig. 3a**). Similarly, pS396-tau^+^ RBPMS^+^ RGC counts were strongly correlated with disease severity ABC scores (**Fig. 3e**; *r*=0.65, P=0.0007), as well as with the cerebral amyloid angiopathy (CAA) grades (**Fig. 3f**; *r*=0.63, P=0.0014). Moderate inverse correlations were detected between GCL Nissl^+^ neuronal % area or RBPMS^+^ RGC counts and CAA grades (**Suppl. Fig. 3b, c**; *r*=-0.42, P=0.06 and *r*=-0.54, P=0.011, respectively).

**Figure 3.**
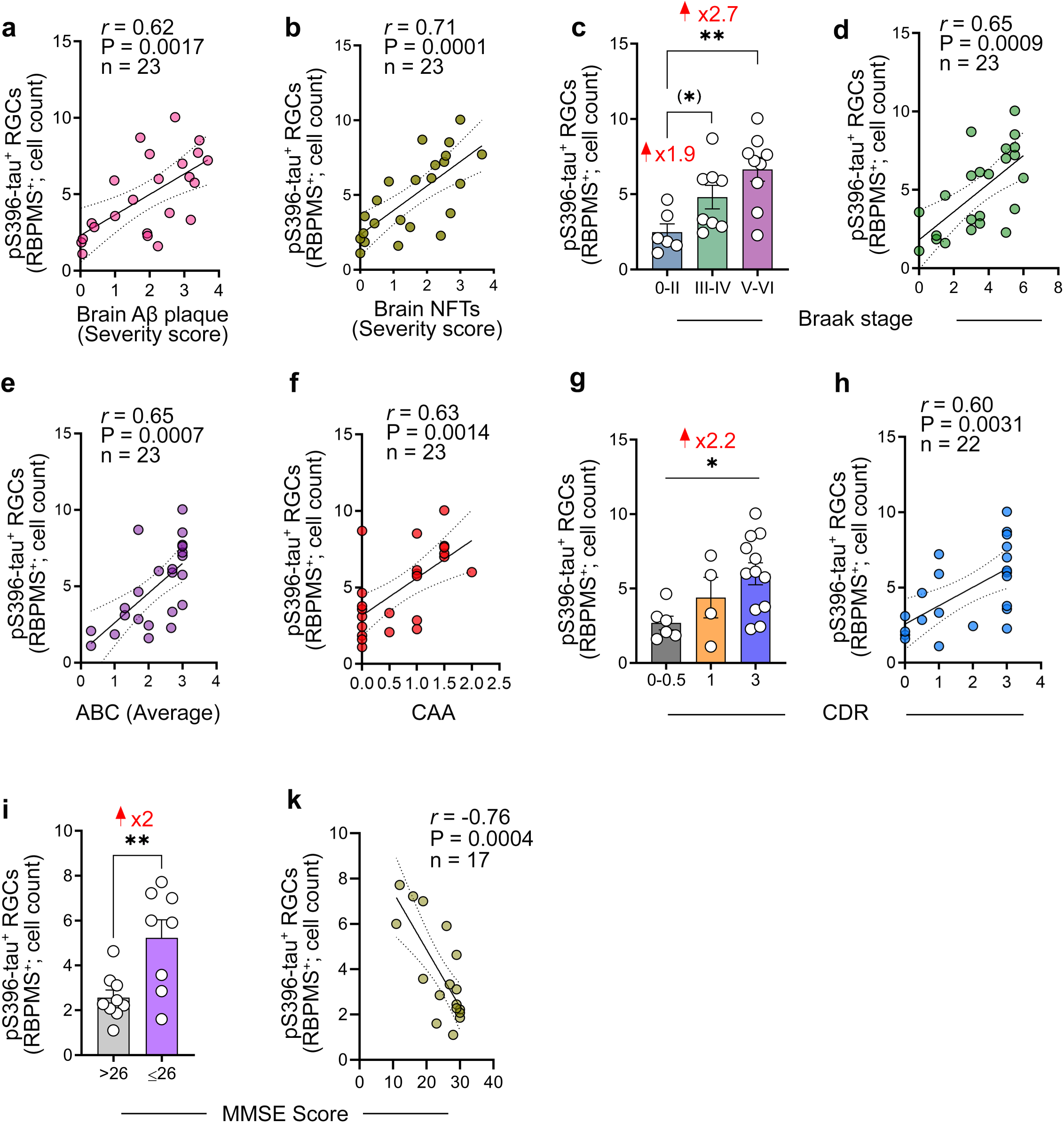
Interactions between retinal pS396-tau in RGCs, brain pathology, and cognitive status. **a, b** Pearson’s correlation coefficient (*r*) analyses between pS396-tau^+^ RGC count and **a** brain amyloid β -protein (Aβ) plaque severity scores, or **b** brain neurofibrillary tangles (NFTs) severity scores. **c** Quantitative analysis of pS396-tau^+^ RGC count stratified by Braak stage classification (n=23) and **d** Pearson’s *r* correlations of pS396-tau^+^ RGC count with the Braak stage. **e, f** Pearson’s correlations between pS396-tau^+^ RGC count and **e** average ABC scores and **f** cerebral amyloid angiopathy (CAA) grade. **g** Quantitative analysis of pS396-tau^+^ RGC count stratified by clinical dementia rating (CDR) scores (n=22) and **h** Pearson’s *r* correlations of pS396-tau^+^ RGC count with the CDR scores. **i** Quantitative analysis of pS396-tau^+^ RGC count stratified by mini-mental state examination score (MMSE) scores (n=17) and **k** Pearson’s *r* correlations of pS396-tau^+^ RGC count with the MMSE scores. Bar graphs are showing individual data points and mean ± SEM. *P < 0.05, **P < 0.01, by one-way ANOVA followed by Tukey’s post-hoc multiple comparison test. Two group comparison is determined by two-tail Student t-test. ABC scores comprise of mean grades for: (A) Aβ plaque score modified from Thal, (B) NFT stage modified from Braak, and (C) neuritic plaque score modified from CERAD.

**Table 2.** Correlations between RGC parameters and brain pathology.

|  | **Aβ (severity score)** |  | **CAA (grade)** |  | **NFT (severity score)** |  | **BRAAK (stage)** |  | **ABC (score)** |  |
| --- | --- | --- | --- | --- | --- | --- | --- | --- | --- | --- |
|  | N = 17-23 |  | N = 17-23 |  | N = 17-23 |  | N = 17-23 |  | N = 17-23 |  |
|  | *r* | P | *r* | P | *r* | P | *r* | P | *r* | P |
| **RBPMs<sup>+</sup> RGCs (count)** |  |  |  |  |  |  |  |  |  |  |
| Total | -0.22 | 0.32 | -0.26 | 0.23 | -0.29 | 0.18 | -0.15 | 0.50 | -0.19 | 0.38 |
| Central | -0.18 | 0.47 | -0.35 | 0.16 | -0.27 | 0.28 | -0.14 | 0.57 | -0.23 | 0.35 |
| Mid-periphery | -0.29 | 0.20 | **-0.54** | **0.011\*** | -0.38 | 0.091 | -0.27 | 0.24 | -0.32 | 0.16 |
| Far-periphery | -0.18 | 0.41 | -0.30 | 0.17 | -0.29 | 0.17 | -0.20 | 0.35 | -0.17 | 0.43 |
| **Nissl<sup>+</sup> in GCL (% area)** |  |  |  |  |  |  |  |  |  |  |
| Total | -0.11 | 0.64 | **-0.42** | **0.060\*** | -0.094 | 0.69 | -0.088 | 0.70 | -0.07 | 0.76 |
| Central | -0.16 | 0.55 | **-0.49** | **0.045\*** | -0.22 | 0.40 | -0.28 | 0.28 | -0.24 | 0.35 |
| Mid-periphery | 0.022 | 0.93 | **-0.48** | **0.028\*** | -0.10 | 0.66 | -0.12 | 0.60 | -0.024 | 0.92 |
| Far-periphery | -0.31 | 0.19 | -0.42 | 0.065 | -0.030 | 0.90 | 0.043 | 0.86 | -0.17 | 0.46 |
| **pS396-tau<sup>+</sup> in RGCs (count)** |  |  |  |  |  |  |  |  |  |  |
| Total | **0.62** | **0.0017\*\*** | **0.63** | **0.0014\*\*** | **0.71** | **0.00010\*\*\*** | **0.65** | **0.0009\*\*\*** | **0.65** | **0.0007\*\*\*** |
| Central | **0.65** | **0.0038\*\*** | **0.59** | **0.010\*** | **0.70** | **0.0014\*\*** | **0.68** | **0.0019\*\*** | **0.67** | **0.0023\*\*** |
| Mid-periphery | **0.51** | **0.017\*** | **0.56** | **0.009\*\*** | **0.72** | **0.00020\*\*\*** | **0.62** | **0.0029\*\*** | **0.57** | **0.0073\*\*** |
| Far-periphery | **0.59** | **0.0029\*\*** | **0.53** | **0.009\*\*** | **0.71** | **0.00020\*\*\*** | **0.63** | **0.0012\*\*** | **0.66** | **0.00070\*\*\*** |

Pearson's correlation analyses: P and *r*-values determine the statistical significance and strength of each pairwise association between retinal RGC marker and brain pathology. P and *r*-values presented in bold type with asterisk(s) are statistically significant (<0.05). Aβ , amyloid beta-protein; CAA, cerebral amyloid angiopathy; NFTs, neurofibrillary tangles. ABC scores comprise of mean grades for: (A) Aβ plaque score modified from Thal, (B) NFT stage modified from Braak, and (C) neuritic plaque score modified from CERAD.

**Table 3.** Correlations between RGC parameters and the cognitive status.

|  | CDR (score) |  | MMSE (score) |  |
| --- | --- | --- | --- | --- |
|  | N = 16-22 |  | N = 15-18 |  |
|  | *r* | P | *r* | P |
| **RBPMs<sup>+</sup> RGCs (count)** |  |  |  |  |
| Total | -0.39 | 0.070 | 0.42 | 0.097 |
| Central | -0.11 | 0.67 | 0.36 | 0.19 |
| Mid-periphery | **-0.45** | **0.047\*** | **0.48** | **0.050** |
| Far- periphery | **-0.49** | **0.022\*** | 0.42 | 0.094 |
| **Nissl<sup>+</sup> in GCL (% area)** |  |  |  |  |
| Total | -0.40 | 0.080 | 0.33 | 0.24 |
| Central | **-0.56** | **0.023\*** | 0.31 | 0.31 |
| Mid-periphery | -0.35 | 0.13 | 0.38 | 0.16 |
| Far-periphery | -0.26 | 0.29 | 0.43 | 0.11 |
| **pS396-tau<sup>+</sup> in RGCs (count)** |  |  |  |  |
| Total | **0.60** | **0.0031\*\*** | **-0.76** | **0.00040\*\*\*** |
| Central | **0.65** | **0.0049\*\*** | **-0.74** | **0.0018\*\*** |
| Mid-periphery | **0.62** | **0.0037\*\*** | **-0.71** | **0.0016\*\*** |
| Far-periphery | **0.61** | **0.0023\*\*** | **-0.59** | **0.0013\*\*** |

Pearson's correlation analyses: P and *r*-values determine the statistical significance and strength of each pairwise association between retinal RGC marker and the cognitive function score. P and *r*-values presented in bold type with asterisk(s) are statistically significant (<0.05). CDR, Clinical Dementia Rating; MMSE, Mini-Mental State Examination.

Finally, we assessed the potential associations between pS396-tau^+^ RBPMS^+^ RGC or RBPMS^+^ RGC counts and cognitive status. Stratification of patients based on their clinical dementia rating (CDR) group revealed a significant 2.2-fold increase in pS396-tau^+^ RGCs in the CDR 3 score group compared to the CDR 0-0.5 group (**Fig. 3g**; P=0.03), with a strong correlation between pS396-tau^+^ RBPMS^+^ RGC count and CDR score (**Fig. 3h**; *r*=0.60, P=0.0031). A moderate correlation was observed between RBPMS^+^ RGC count and CDR score (**Suppl. Fig. 3d;** *r*=-0.49 and P=0.022). Importantly, stratifying patients based on the mini-mental state examination (MMSE) cut-off score of 26, which has been reported to have high sensitivity and specificity for detecting dementia [87], was utilized in our cohort. This analysis showed a significant 2-fold increase in pS396-tau^+^ RBPMS^+^ RGC counts in the MMSE ≤ 26 group compared to the MMSE >26 group (**Fig. 3i**; P=0.0059). Whereas no significant association was detected between RBPMS^+^ RGC counts and MMSE score (**Suppl. Fig. 2e**), a highly significant and strong association was observed between pS396-tau^+^ RBPMS^+^ RGC count and MMSE score (**Fig. 3k**; *r*=-0.76, P=0.0004).

## Discussion

In this study, we present the first evidence of abnormal tau inclusions within RBPMS-positive RGCs, concomitant with ganglion cell loss in the retinas of donor patients with MCI (due to AD) and AD dementia. Increases in pS396-tau-containing RBPMS-positive RGCs in both MCI and AD patients were accompanied by elevated apoptotic cell markers, necroptotic-like morphological changes in RGCs, including hypertrophic soma and nuclei displacement, and decreased RGC counts. Tau oligomers were also detected in swollen RGCs within the GCL. Notably, we found moderate to strong associations between RGC loss and pS396-tau burden in both RGCs and the retina as a whole. Moreover, we observed that retinal tau oligomers, as well as retinal Aβ_42_ and intra-RGC Aβ oligomers, were strongly associated with RGC reduction, suggesting a link between retinal tau and amyloid pathologies and ganglion cell degeneration in AD. Importantly, our data indicated tight correlations between pS396-tau-containing RBPMS^+^ RGCs and the respective brain pathology, disease stage, and cognitive status. Overall, our findings suggest that abnormal tau isoforms accumulate within RBPMS^+^ RGCs and are associated with early and marked RGC loss in AD patients.

Among the RGC populations, the midget cells projecting to the parvocellular (P-cell) layers of the lateral geniculate nucleus (LGN) and the parasol cells projecting to the magnocellular (M-cell) layers of the LGN serve as two distinct visual pathways that process color and low spatial frequency contrast vision, respectively [69]. In AD patients, abnormalities in color vision, eye movement, contrast sensitivity, and visual integration have been detected early in disease progression [37, 47, 78] [32, 72, 101]. Therefore, fluctuations in color perception and abnormal contrast sensitivity in AD patients may be attributed to damage and loss of these RGC types, in addition to the involvement of horizontal and amacrine neurons. Here, the analysis of the RBPMS marker, a conserved RNA binding protein with a single RNA recognition motif expressed in RGCs of humans and animal models [84, 91], facilitates differentiation from other retinal cells [63, 92, 98] and further validates our findings in RGCs. Notably, our analysis of RGC integrity in the superior temporal retina indicates marked decreases in RBPMS^+^ RGC counts or immunoreactive area (by 47-55% in MCI and 46-50% in AD) compared to CN controls, with similar degrees of decreases observed in GCL Nissl^+^ neurons (by 56% in MCI and 55% in AD patients). These results are consistent with previous studies reporting significant reductions in RGCs and the GCL in AD patients versus control subjects [6, 15–17] [101].

Specifically, the RBPMS^+^ RGC count per retinal subregion indicated a 46% loss in MCI and a 57% loss in AD in the mid-periphery, as well as a 62% loss in MCI and a 45% loss in AD in the far periphery. Similarly, a study by Blanks et al., described GCL neuronal loss in AD as most pronounced in the superior and inferior quadrants, ranging between 40% and 49% throughout the mid-peripheral subregions and reaching 50-59% in the far-peripheral retina of AD patients [16]. These peripheral retinal subregions, which have anatomically fewer ganglion cells and a thinner nerve fiber layer, appear more vulnerable to RGC loss in AD, potentially due to a higher density of abnormal Aβ and tau species (e.g., Aβ_42_, Aβ oligomers, PHF-tau, pS396-tau and p-tau (S202/T205)), and microgliosis [60, 61, 70, 108]. Interestingly, whereas the total and mid-peripheral ST retina consistently demonstrated significant and similar RGC reductions in both MCI and AD patients, as shown by the GCL Nissl^+^ area and RBPMS^+^ RGC count analyses, non-significant trends were noted for the far and central subregions, respectively. These differences may be due to variations in the types of analysis and staining patterns. The loss of RGCs in MCI and AD patients may explain previous reports of visual dysfunctions in AD, specifically impaired color and low spatial frequency contrast vision, as well as motion perception that can be attributed, at least in part, to the loss of M-cell and P-cell RGCs. In addition, a previous study of postmortem AD retinas identified a reduction in melanopsin retinal ganglion cells (mRGCs), intrinsically photosensitive cells that contribute to the photoentrainment of circadian rhythms, potentially explaining the sleep disturbances observed in these patients [67].

In the brains of AD patients, the increase in hyperphosphorylated tau isoforms has been shown to lead to tau aggregation, oligomerization, propagation, and NFT formation, ultimately causing neuronal dysfunction and degeneration [90, 114]. Previous studies detected intracellular pretangles and mature tangles in the retinas of AD patients [27, 29, 40, 45, 46, 60, 86, 108, 122]. Recently, we also identified tau oligomers and citrullinated-tau, along with other tau isoforms, in the retinas of MCI and AD patients. Notably, both pS396-tau and oligomeric-tau forms were frequently observed within the GCL, with significant increases in MCI and AD patients [108]. The pS396-tau isoform is increased in the AD brains and is linked to neuronal cell loss and Braak stage severity [8, 36, 96, 120]. Here, we found a specific build-up of these pathological tau isoforms within RGCs of MCI and AD patients, demonstrating their connection with RGC integrity, entailing similar links to neurodegeneration and tauopathy as seen in the brain.

In this study, we detected higher numbers of RBPMS^+^ RGCs containing pS396-tau in patients with AD dementia and those at the earliest stages of functional impairment (MCI due to AD). The level of pS396-tau in RGCs was even higher in AD patients compared to MCI patients, suggesting that more RGCs are affected by pS396-tau as the disease progresses. Our data on pS396-tau^+^ RGC counts per retinal subregion indicated that the most significant and substantial changes were detected in the central subregion across all analyzed groups, with less pronounced changes in the mid-periphery and the far periphery. This could be attributed to the density of RGCs in each subregion, as there are up to eight layers of ganglion cells in the central subregion and only one or two layers with space between them in the far periphery [55]. Hence, there is a higher probability that RGCs in the central subregion are impacted by pS396-tau compared to those in the peripheral subregions. These findings may guide potential future in vivo imaging of pS396-tau-positive RGCs in the central ST retina for early AD detection and monitoring of disease progression.

In both fluorescent and peroxidase-based staining methods, we observed a three-parallel-string staining pattern of retinal pS396-tau in the IPL of MCI and AD patients. Consistent with retinal neuroanatomy, these findings suggest that pS396-tau accumulates within the neuronal dendrites of RGCs, which connect with the axons of bipolar and amacrine cells. These tau aggregates in synaptic-rich regions may interfere with information transmission and could help explain the decrease in contrast sensitivity observed in MCI and AD patients. Moreover, in MCI and AD patients, the pS396-tau isoform was also observed in the OPL, specifically in horizontal cells. A recent study suggested that pS202/T205-tau (AT8^+^) spreads from the OPL to the IPL/GCL in the AD retina [122]. The patterns of retinal pS396-tau burden in the NFL of CN subjects and the IPL/OPL of MCI and AD patients merit further investigation to understand how pS396-tau spreads across retinal layers and neuronal processes during AD progression.

Our analysis showed a moderate inverse correlation between pS396-tau^+^ RGCs and RGC integrity, and a stronger negative correlation between overall retinal pS396-tau burden and RGC integrity, suggesting that the extent of retinal pS396-tau load, including in neuronal dendrites connecting with RGCs, may have additive effects on RGC susceptibility. Beyond retinal p-tau, the strong negative associations of Aβ oligomers in RGCs and retinal tau oligomers with RGC reduction suggest their substantial and detrimental effects on RGC degeneration. These retinal findings in AD are consistent with similar reports connecting elevated Aβ and tau oligomers with neuronal loss in AD brains [3, 9, 68, 83, 105, 116]. As expected, the overall burden of retinal pS396-tau strongly correlated with the extent of pS396-tau-loaded RGCs. Unexpectedly, the levels of retinal Aβ_42_ also strongly correlated with the extent of RGC containing pS396-tau, suggesting that retinal Aβ may be a driver of tauopathy in RGCs, similar to the interactions between Aβ and the spread of tau in neurons of AD brains [18, 126].

Levels of the early apoptotic marker, cleaved caspase 3 [117], have been shown to be elevated in AD brains, with a high degree of colocalization to neurofibrillary tangles within neurons [38, 112]. In the current study, we observed increased cleaved caspase 3 expression in GCL cells in AD, but not MCI, patients compared to cognitively normal controls. This is consistent with previous studies showing cleaved caspase 3^+^/Tuj1^+^ RGCs [40] and overall retinal cleaved caspase 3 expression [61] in AD patients compared to controls. The elevated expression of retinal cleaved caspase 3 in GCL cells, along with strong correlations with pS396-tau-loaded RGCs, suggests that pS396-tau may trigger apoptotic cell death in RGCs.

RGCs are highly diverse, consisting of multiple subtypes that exhibit a range of morphological and physiological characteristics, including variations in soma and cell body size [55]. In this study, we observed that RGCs in aged CN individuals predominantly appear to have small-sized somas, with a minority of cells exhibiting large and round somas. In contrast, a substantial number of RGCs in MCI and AD patients appeared swollen, with enlarged somas, granulovacuolar-like bodies, and displaced nuclei, particularly in those containing pS396-tau inclusions. To the best of our knowledge, this is the first demonstration of hypertrophic RGCs in MCI and AD patients. This abnormal RGC morphology is characteristic of neurons exhibiting granulovacuolar degeneration due to necroptosis, a process observed in the brains of individuals with preclinical AD and AD dementia [59]. The morphology and process of necroptotic cells are characterized by compromised plasma membrane integrity, organelle and cell enlargement, chromatin fragmentation, and eventual cell lysis [89, 125]. Moreover, studies have indicated that necroptosis is involved in AD brain pathology and is closely linked to tau pathology and Braak stage progression [11, 21, 59], with recent research showing that p-tau contributes to neuronal death by inducing necroptosis and inflammation [28]. In this study, the abnormal morphology of RGCs, particularly in those with a pS396-tau burden, may indicate necroptotic cell death in the RGCs of AD retinas. Future studies are needed to determine the potential role of p-tau in retinal ganglion cell death in AD.

Looking into the potential connections between pS396-tau-containing RGCs and disease status, our analysis indicates strong associations between pS396-tau^+^ RGCs and the following brain pathologies: Aβ plaques, NFTs, Braak stage, and ABC neuropathic changes. However, RGC counts alone did not correlate with these AD brain parameters. These data suggest that tauopathy-laden RGCs (measured here by RBPMS^+^ RGCs with a pS396-tau burden) may represent the link between retinal neuronal injury and brain AD pathology and disease progression. The strong correlation between pS396-tau-positive RGCs and CAA severity may simply reflect brain Aβ burden, as CAA involves cerebrovascular deposition of Aβ and is influenced by Aβ plaque levels [39]. In our cohort, pS396-tau-containing RGCs had comparable correlations with brain Aβ plaques and CAA severity. Importantly, our data indicate that pS396-tau-containing RGC numbers strongly correlate with cognitive status, as measured by the CDR, and even more so with MMSE scores. While the CDR is a test that allows assessment of cognitive, behavioral, and functional performance associated with AD, the MMSE test evaluates cerebral competency, comprehension, and communication. Our findings suggest that a future retinal imaging approach that reliably measures the number of RGCs containing pS396-tau in the ST central region holds potential as a marker to evaluate brain NFT severity, Braak staging, ABC scores, and cognitive deficits in AD patients. In the clinical setting, GCL layer is assessed by OCT [37, 73, 75, 109], apoptotic RGCs can be images by detection of apoptosing retinal cells (DARK) method [10, 26], and more specific RGC changes could be detectable in the inner retina using high resolution imaging systems, such as AO-OCT. Imaging RGCs, combined with future p-tau tracers, could serve as a non-invasive biomarker for early AD diagnosis and monitoring of disease progression. This would be immensely valuable in future trials evaluating new treatments for AD.

We acknowledge several limitations of this study. As a cross-sectional, case-control study, our focus was primarily on group stratification and correlations, so caution must be exercised before implicating cause-and-effect conclusions. Moreover, the lack of clinical information on visual system-related symptoms hinders our ability to assess potential connections between pS396-tau^+^ RBPMS^+^ RGCs and various manifestations of visual dysfunction. This highlights the need for future studies to explore the relationships between pS396-tau^+^ RGCs, RGC loss, and ocular outcomes in patients. Future studies in larger and more diverse populations are warranted to validate these findings and to compare RGC susceptibility with that of other retinal cell types in relation to AD processes.

## Conclusion

In summary, this study provides the first evidence of RGCs laden with abnormal tau inclusions, pS396-tau and oligomeric tau, in early (MCI) and advanced-stage AD patients, with clear indications of increased RGC vulnerability. RBPMS-positive RGCs containing pS396-tau correlated with increased apoptotic markers, necroptotic-like morphological changes, and reduced RGC counts, suggesting that these tau pathologies may contribute to ganglion cell degeneration in AD. Notably, strong correlations were found between pS396-tau laden RGCs and brain AD pathology, cognitive status, and disease stage. This study highlights the potential of imaging tau-laden RGCs as a non-invasive biomarker for early AD diagnosis and monitoring disease progression. However, further research is needed to more definitively establish these connections.

## Supporting information

Davis et al_Supplementary material

## Abbreviations

A: Amyloid
Ab: Antibody
ABC: Amyloid/Braak/CERAD score
AD: Alzheimer’s disease
AD: RCAlzheimer’s disease research center
Aβ: Amyloid β-protein
ANOVA: Analysis of variance
B: Brain
C: Central retina
CCasp3: Cleaved caspase 3
CDR: Clinical Dementia Rating
CN: Cognitively normal
F: Far-peripheral retina
GCL: Ganglion cell layer
IHC: Immunohistochemistry
INL: Inner nuclear layer
IPL: Inner plexiform layer
IR area: Immunoreactive area
mAb: Monoclonal antibody
M: Middle-peripheral retina
MCI: Mild Cognitive Impairment
MMSE: Mini-mental state examination
mRGC: Melanopsin Retinal Ganglion Cell
NDRI: National disease research interchange
NFL: Nerve fiber layer
NFT: Neurofibrillary tangle
NT: Neuropil thread
OD: Optic disc
ONL: Outer nuclear layer
OPL: Outer plexiform layer
pAb: Polyclonal antibody
PMI: Postmortem interval
p-tau: Hyperphosphorylated tau
RBPMS: Ribonucleic acid binding protein with multiple splicing
RGC: Retinal Ganglion Cell(s)
Serine 396: S396
SD: Standard deviation
ST: Superior temporal.

## Declarations

## Acknowledgments

We thank Elijiah Maxfield for assisting with manuscript editing. We thank Prof. Carol Ann Miller, the former director of the USC-ADRC neuropathology core laboratory, for providing the neuropathological reports. We thank the late Prof. Peter Davies of The Litwin-Zucker Research Center for the Study of Alzheimer’s Disease, The Feinstein Institutes for Medical Research, New York, for generously providing the PHF-1 antibodies. We thank Profs. Giovanni Meli and Antonino Cattaneo for providing antibodies against Aβ oligomers (scFvA13). The authors dedicate this manuscript to the memory of Dr. Salomon Moni Hamaoui and Lillian Jones Black, both of whom died from Alzheimer’s disease.

## Ethic approval and Consent to participate

This study is not considered a human subjects research, and we confirm that consent was not necessary, for the reasons described as follow: we processed and analyzed deidentified retinal tissues of deceased patients that were provided by the USC-ADRC (IRB protocol HS-042071) and NDRI (Cedars-Sinai Medical Center IRB protocol Pro00019393). Histological studies at Cedars-Sinai Medical Center were performed under IRB protocols Pro00053412 and Pro00019393.

## Author contributions

M.R.D.: performed experiments, collected, and analyzed data, created figures, drafted and edited the manuscript. E.R., A.R., B.P.G., E.S.-G.: performed experiments, collected, and analyzed data. D.-T.F.: analyzed data, created figures and illustrations, edited the manuscript. Y.K.: performed experiments, analyzed data, assisted in creating figures, wrote and edited the manuscript. N.M.: assisted with experimental design and execution, collected data. L.S., D.H. provided donor eyes and the clinical and brain pathological data. R.K.: provided the T22 antibodies that recognize tau oligomers. A.A.S., A.V.L, K.L.B.: assisted with interpretation of data and editing. M.K.-H. was responsible for study conception and design, data analysis and collection, interpretation of data, study supervision, and manuscript writing and editing. All authors have read and approved the manuscript.

## Consent for publication section

Not applicable

## Availability of data and material

Data generated and analyzed in this study are included in this published manuscript and the supplementary online material. Additional data can be made available by the corresponding PI upon reasonable request.

