## Supplementary material for "Retinal ganglion cell vulnerability to pathogenic tau in Alzheimer’s disease": Davis et al_Supplementary material

**Supplementary Table 1**. List of human donors in this study.

**Supplementary Table 2.** List of antibodies.

**Supplementary Figure 1.** Additional data on RGC loss in MCI and AD patients.

**Supplementary Figure 2.** Additional data on pS396-tau within RGCs of MCI and AD patients.

**Supplementary Figure 3.** Additional correlation analyses.

**Supplementary Table 1. List of human donors in this study.**

| Diagnosis | Sex | Race | Age at death | Thal A | Braak B | CERAD C | CAA Score | Co-morbid. [LB/AS VD] | Braak Stage | CDR Score | MMSE Score | APOE status |
|---|---|---|---|---|---|---|---|---|---|---|---|---|
| AD1 | M | W | 79 | 3 | 3 | 3 | 1.5 | -/+ | V | n.a. | n.a. | n.a. |
| AD2 | M | H | 97 | 3 | 2 | 3 | 1 | -/+ | III | 1 | 26 | e3/e3 |
| AD3 | M | W | 90 | 3 | 3 | 3 | 2 | -/+ | VI | 3 | n.a. | n.a. |
| AD4 | M | W | 90 | 3 | 2 | 3 | 1 | -/+ | IV | 3 | n.a. | e3/e4 |
| AD5 | F | W | 90 | 3 | 3 | 3 | 1 | -/+ | V-VI | 3 | n.a. | e3/e4 |
| AD6 | F | H | 81 | 3 | 3 | 3 | 1.5 | -/+ | V-VI | 3 | 12 | e3/e3 |
| AD7 | F | W | 94 | 3 | 3 | 3 | 0 | -/+ | V-VI | 3 | n.a. | e3/e3 |
| AD8 | F | W | 85 | 3 | 3 | 3 | 1.5 | -/+ | V-VI | 3 | n.a. | e3/e3 |
| AD9 | M | W | 66 | 3 | 3 | 3 | 1.5 | -/+ | V | 3 | 19 | n.a. |
| AD10 | M | W | 88 | 3 | 3 | 3 | 1.5 | -/+ | V-VI | 1 | 16 | e3/e2 |
| AD11 | F | W | 86 | 3 | 3 | 2 | 1 | -/+ | V-VI | 3 | 18 | e3/e4 |
| AD12 | F | W | 90 | 2 | 3 | 3 | 1 | -/+ | V | 2 | 9 | n.a. |
| AD13 | F | W | 100 | 2 | 3 | 3 | 0.5 | -/+ | V-VI | 2 | 16 | n.a. |
| AD14 | M | W | 88 | 2 | 3 | 2 | 1 | -/+ | V-VI | 1 | 4 | e3/e4 |
| AD15 | F | A | 88 | 2 | 3 | 3 | 1.5 | -/+ | V | 3 | 4 | n.a. |
| MCI1 | F | W | 89 | 1 | 2 | 2 | 1 | -/+ | III-IV | 0.5 | 24 | e3/e3 |
| MCI2 | M | H | 80 | 3 | 3 | 2 | 1 | -/+ | V | 3 | 29 | e3/e3 |
| MCI3 | M | W | 93 | 2 | 0 | 2 | 0 | -/+ | 0 | 3 | 19 | e3/e2 |
| MCI4 | F | W | 93 | 3 | 2 | 2 | 2 | -/+ | IV | 3 | 11 | e3/e3 |
| MCI5 | F | W | 87 | 3 | 3 | 3 | 1.5 | -/+ | V-VI | 3 | 13 | e3/e3 |
| MCI6 | F | W | 86 | 3 | 1 | 3 | 0 | -/- | I-II | 2 | 15 | e3/e4 |
| MCI7 | F | W | 76 | 3 | 3 | 3 | 2 | -/+ | V | 1 | 26 | e3/e4 |
| MCI8 | F | W | 98 | 2 | 3 | 2 | 2 | -/- | V | 2 | 15 | n.a. |
| MCI9 | M | W | 88 | 1 | 2 | 2 | 0 | -/+ | III | 3 | n.a. | n.a. |
| MCI10 | F | B | 94 | 2 | 1 | 2 | 0 | -/- | I-II | 0.5 | 29 | e3/e3 |
| CN1 | M | W | 95 | 1 | 1 | 1 | 0 | -/- | I | 0 | 30 | e3/e3 |
| CN2 | F | W | 99 | 1 | 2 | 1 | 0 | -/- | III | 0 | 29 | e3/e3 |
| CN3 | M | H | 81 | 3 | 1 | 2 | 0.25 | -/- | I-II | 0 | 23 | e3/e4 |
| CN4 | F | W | 93 | 3 | 2 | 3 | 0.5 | -/+ | III-IV | 1 | 27 | e3/e2 |
| CN5 | F | W | 66 | n.a. | n.a. | n.a. | n.a. | n.a. | n.a. | n.a. | n.a. | n.a. |
| CN6 | M | W | 84 | n.a. | n.a. | n.a. | n.a. | n.a. | n.a. | n.a. | 30 | n.a. |
| CN7 | F | B | 73 | n.a. | n.a. | n.a. | n.a. | n.a. | n.a. | n.a. | n.a. | n.a. |
| CN8 | F | W | 58 | n.a. | n.a. | n.a. | n.a. | n.a. | n.a. | n.a. | 30 | n.a. |
| CN9 | F | W | 76 | n.a. | n.a. | n.a. | n.a. | n.a. | n.a. | n.a. | n.a. | n.a. |
| CN10 | F | W | 75 | n.a. | n.a. | n.a. | n.a. | n.a. | n.a. | n.a. | n.a. | n.a. |
| CN11 | M | W | 70 | n.a. | n.a. | n.a. | n.a. | n.a. | n.a. | n.a. | 30 | n.a. |
| CN12 | F | W | 86 | n.a. | n.a. | n.a. | n.a. | n.a. | n.a. | n.a. | n.a. | n.a. |

| Diagnosis | Sex | Race | Age at death | Thal A | Braak B | CERAD C | CAA Score | Co-morbid. [LB/ASVD] | Braak Stage | CDR Score | MMSE Score | APOE status |
|---|---|---|---|---|---|---|---|---|---|---|---|---|
| CN13 | M | W | 77 | n.a. | n.a. | n.a. | n.a. | n.a. | n.a. | n.a. | 30 | n.a. |
| CN14 | F | W | 95 | 1 | 0 | 0 | 0.5 | -/- | I | 0 | 30 | n.a. |
| CN15 | M | W | 69 | 0 | 0 | 1 | 0 | -/+ | 0 | 1 | 28 | n.a. |
| CN16 | F | W | 91 | 2 | 2 | 2 | 0 | -/+ | III | 2 | 29 | n.a. |

AD, Alzheimer's disease dementia; MCI, mild cognitive impairment; CN, normal cognition; F, female; M, male; A, Asian; B, Black; H, Hispanic; W, White; A, Aβ plaque score modified from Thal; B, NFT stage modified from Braak; C, Neuritic plaque score modified from CERAD; CAA, Cerebral amyloid angiopathy; LB, Lewy bodies; ASVD. Atherosclerosis; CDR, Clinical dementia rating; MMSE, Mini-Mental State Examination; n.a., not available; +: present; –: none; APOE, apolipoprotein alleles.

**Supplementary Table 2.** List of antibodies for immunohistochemistry.

| ***Primary antibody*** | **Source Species** | **Dilution** | **Application** | **Commercial Source** | **Catalog Number** |
|---|---|---|---|---|---|
| RBPMS pAb | Rabbit | 1:300 | IF | GeneTex | GTX118619 |
| Parvalbumin pAb | Goat | 1:200 | IF | Novus Biologicals | NB100-1541 |
| CCasp3 pAb | Rabbit | 1:400 | IF | Cell Signal | 9661 |
| VGLUT1 pAb | Guinea Pig | 1:1000 | IF | Chemicon | AB5905 |
| Ser396 (pS396-tau) pAb | Rabbit | 1:1200 | IF, DAB | Anaspec | AS-54977 |
| T22 (oligo-tau) pAb | Rabbit | 1:200 | IF | Prof. Rakez Kayed Lab | - |
| PHF-1-tau mAb | Mouse | 1:200 | IF | Prof. Peter Davies Lab | - |
| scFvA13 (Oligo-Aβ) mAb | Mouse* | 1:450 | IF | Prof. Giovanni Meli Lab | - |
| 12F4 ($A\beta_{42}$) mAb | Mouse | 1:500 | IF | Biolegend | 805502 |
| CCasp3 pAb | Rabbit | 1:250 | IF | Cell signaling | 9661 |
| ***Secondary antibody*** | | | | | |
| Cy2 (anti-Goat, Mouse, Rabbit) | Donkey | 1:200 | IF | Jackson ImmunoResearch Laboratories | |
| Cy3 (anti-Goat, Mouse, Rabbit, Guinea Pig ) | Donkey | 1:200 | IF | | |
| Cy5 (anti-Goat, Mouse, Rabbit, Guinea Pig) | Donkey | 1:200 | IF | | |

Abbreviations: IF – immunofluorescence; DAB – peroxidase-based immunohistochemistry visualized with DAB (3, 3'-diaminobenzidine) substrate; Cyanine dyes – Cy2, Cy3, Cy5; pAb – polyclonal antibody; mAb – monoclonal antibody; p-tau – hyperphosphorylated tau; oligo-tau – oligomeric tau forms; PHF – paired-helical filament (pS396/pS404); scFv – single chain Fv fragment VGLUT1 – Vesicular glutamate transporter 1; *mouse recombinant antibody fragment.

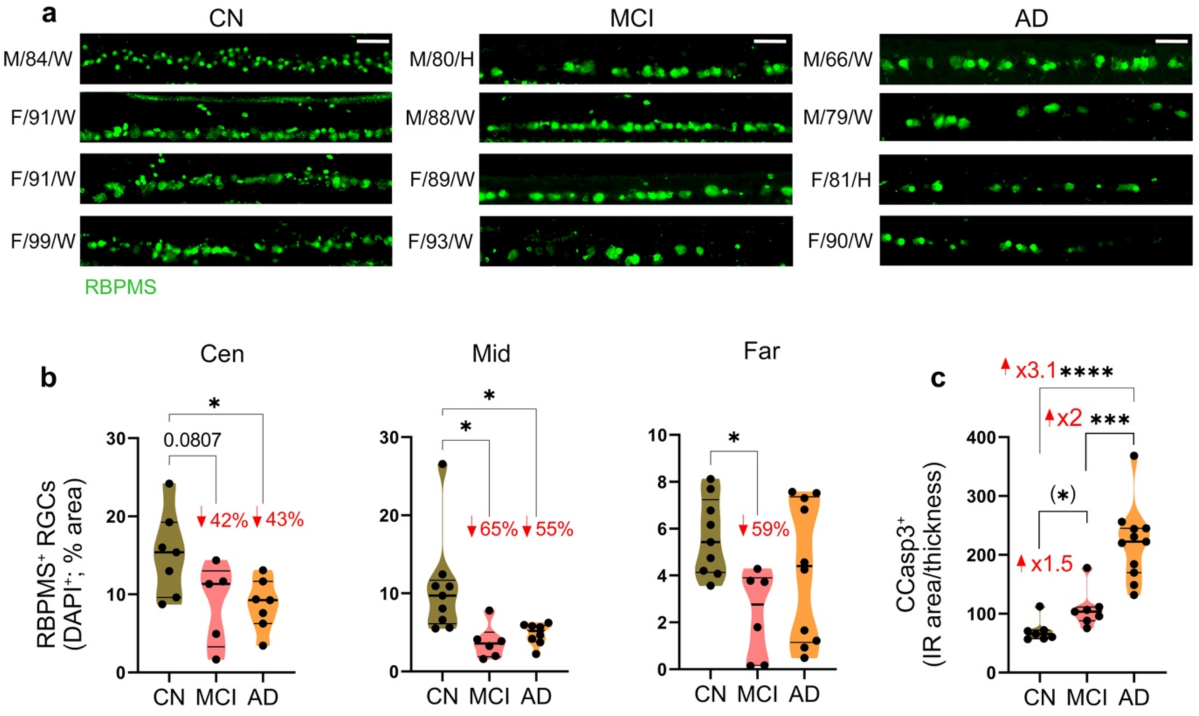


**Supplemental Figure 1.** Extended data on ganglion cell integrity in retinal tissues from MCI and AD patients.

**a** Representative microscopic images of RBPMS$^+$ RGCs within the GCL, labeled with RBPMS (green), in retinal cross-sections from patients with mild cognitive impairment (MCI due to AD, n=4) and Alzheimer's disease (AD) dementia (n=4), compared to cognitively normal individuals (CN, n=4) Scale bar: 50 µm. **b** Quantitative immunohistochemistry analysis of RBPMS$^+$DAPI$^+$ RGCs % area in Cen, Mid-, and Far-peripheral ST subregions (n=19-25 subjects). **c** Total Cleaved Caspase-3 (CCasp3)$^+$ immunoreactive area analysis normalized to retinal thickness (n=25 subjects; n=7 CN, n=7 MCI, n=11 AD). Individual data points (circles) and median, lower and upper quartile are shown in violin plots. *P < 0.05, ***P < 0.001, ****P < 0.0001, by one-way ANOVA with Tukey's post-hoc multiple comparison test or two-tailed Student t-test (in parenthesis). Percent decreases and fold changes are shown in red. F, Female; M, Male; Age (in years); Ethnicity: W, White and H, Hispanic.

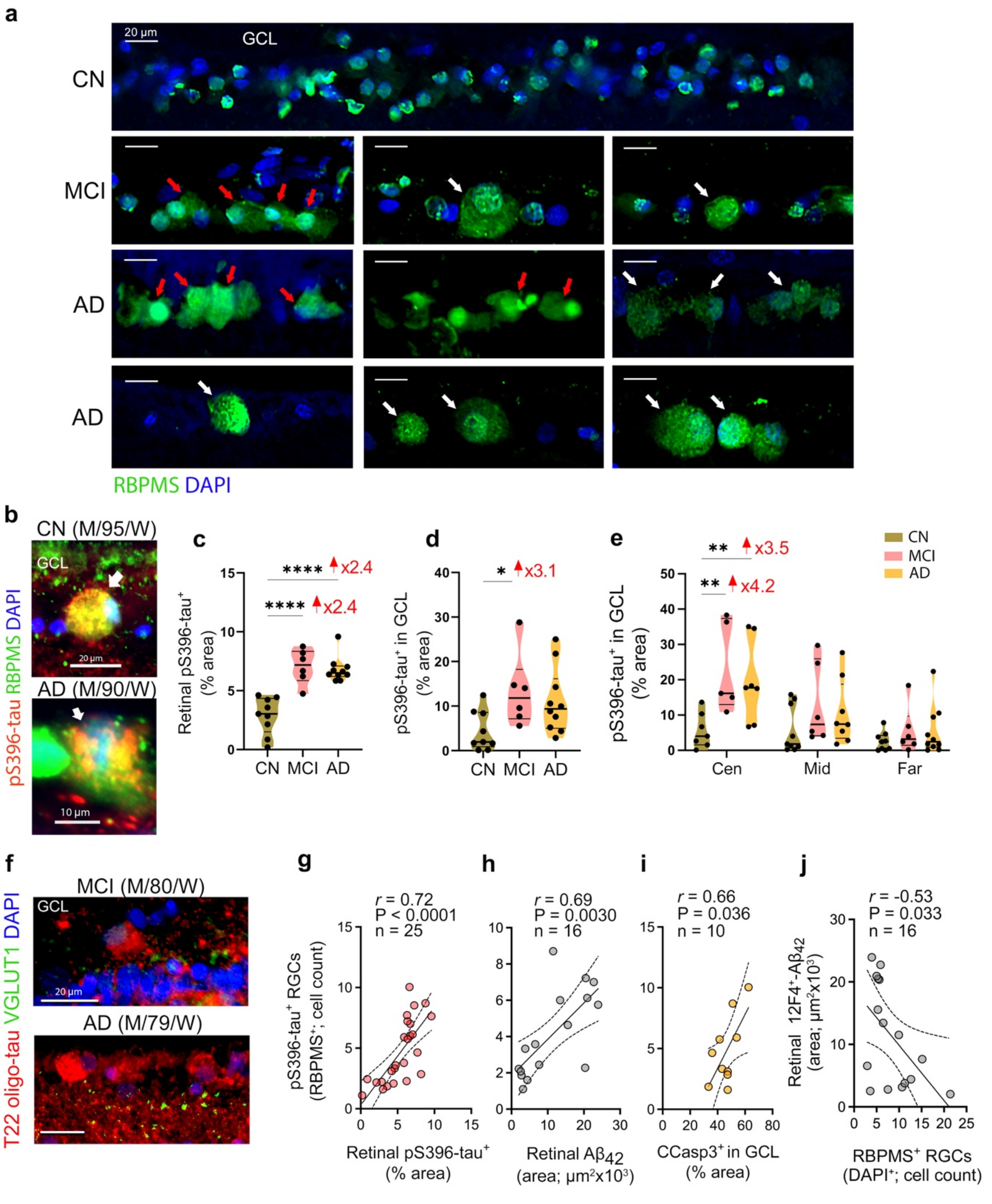


**Supplemental Figure 2.** Extended data on RGC morphology and pS396-tau within RGCs in MCI and AD patients.

**a** Representative microscopic images of retinal cross-section immunofluorescently stained for $RBPMS^+$ showing abnormal morphology of RGCs in MCI and AD retinas as compared with CN controls. Red arrows point to nuclear displacement and white arrows indicate granulomatous cytoplasm. **b** High-magnification microscopic images depicting pS396-tau accumulation (red) in swollen $RBPMS^+$ RGCs (green) with hypertrophic soma (white arrows) in CN and AD retinas. **c, d** Quantitative immunohistochemistry analysis of **c** retinal pS396-tau$^+$ % area and **d** pS396-tau$^+$ % area in the GCL (n=25 subjects; n=9 CN, n=6 MCI, n=10 AD). **e** Analyses of pS396-tau % area in the GCL per Cen, Mid- and Far-peripheral ST subregions (n=19-25). **f** Immunostaining for $T22^+$ oligo-tau (red) in the GCL, together with post-synaptic marker, VGLUT1 (green), and DAPI for nuclei (blue) in MCI and AD retinas. Scale bars: 20 µm and 10 µm. **g-i** Pearson's correlation (*r*) analyses between pS396-tau$^+$ RGCs count and **f** retinal pS396-tau$^+$ % area, **h** retinal 12F4$^+$A$\beta_{42}$ immunoreactive area and **i** CCasp3$^+$ % area in GCL. **j**. Pearson's correlation analyses between retinal 12F4$^+$A$\beta_{42}$ and RBPMS$^+$DAPI$^+$ RGCs. Individual data points (circles) and median, lower and upper quartile are shown in violin plots. *P < 0.05, **P < 0.01, ****P < 0.0001, by one-way or two-way ANOVA with Tukey's post-hoc multiple comparison test. Fold changes are shown in red. F, Female; M, Male; Age (in years); Ethnicity: W, White.

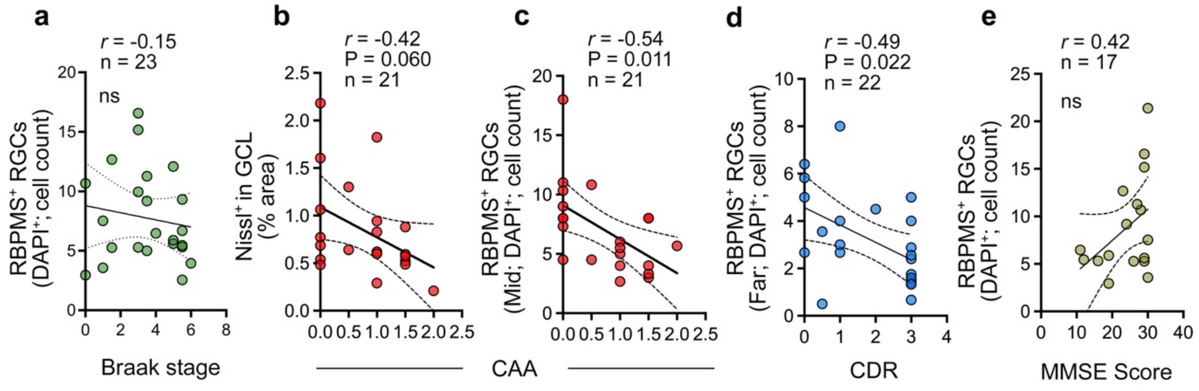


**Supplemental Figure 3.** Extended correlations between retinal RGC markers and brain pathology and cognitive status.

**a-e** Pearson's correlation coefficient (*r*) analyses between **a** RBPMS$^+$ RGCs count and Braak stage, **b** Nissl$^+$ % area in GCL and CAA grade, **c** RBPMS$^+$ RGC count in mid-periphery and CAA grade, **d** RBPMS$^+$ RGC count in far-periphery and the CDR score, and **e** RBPMS$^+$ RGC total count and MMSE score. CDR, Clinical Dementia Rating; MMSE, Mini-mental state examination; RGCs, retinal ganglion cells; RBPMS, Ribonucleic acid binding protein with multiple splicing.
